# Dengue Virus NS1 Binds Ephrin B1 to Trigger Endothelial Dysfunction

**DOI:** 10.1101/2025.11.19.689067

**Authors:** Felix Pahmeier, Sabrina R. Hammond, Charlotte Flory, Xinyi Feng, Saeyoung E. Lee, Erika V. Jimenez-Posada, Elias M. Duarte, Jaime A. Cardona-Ospina, Aquena H. Ball, Nharae E. Lee, Kayla Leung, Laurentia V. Tjang, P. Robert Beatty, Scott B. Biering, Pietro Scaturro, Eva Harris

## Abstract

Dengue virus (DENV), a member of the *Flaviviridae* family, is the most prevalent and medically important mosquito-borne viral pathogen. Infected cells secrete the viral non-structural protein 1 (NS1) into the bloodstream, where it can interact with endothelial cells to induce vascular leak. Host factors involved in NS1-mediated endothelial dysfunction are incompletely understood, leading us to investigate the host interactome of DENV NS1 in endothelial cells via a comparative mass spectrometry approach. We identified ephrin B1 (EFNB1) as a critical host factor in NS1-mediated endothelial barrier dysfunction and show that phosphorylation of EFNB1 is necessary for induction of barrier dysfunction. Further, we map the interface of the EFNB1-NS1 complex through biochemical and computational approaches, and we show that EFNB1-Fc-fusion proteins can act as decoys to block NS1-induced barrier dysfunction *in vitro* and *in vivo*. This study provides insights into the mechanism of flavivirus NS1-mediated endothelial barrier dysfunction and new avenues to target vascular leak.

## Main text

Mosquito-borne flaviviruses are medically important viruses that pose a major global public health threat, with more than half of the world’s population at risk of infection^1,2^. Members include the four dengue virus serotypes (DENV1-4), Zika virus (ZIKV), West Nile virus (WNV), and yellow fever virus (YFV). Case numbers have been increasing in recent years and continue to rise due to global trade and travel, climate change, and expansion of the geographic range of the *Aedes* mosquito vectors^1^. Vaccine development has been difficult, particularly for DENV and ZIKV, due to the potential for antibody-dependent enhancement (ADE) of cross-reactive antibodies elicited by natural infection and vaccines^3–5^. Further, there are currently no FDA-approved treatment options available.

Flaviviruses encode a single-stranded, positive-sense RNA genome encoding three structural and seven non-structural proteins. The non-structural protein 1 (NS1) is secreted from infected cells as an oligomer (tetramer or hexamer) that contains a lipid cargo^6–9^. In addition to antibodies targeting the structural proteins, flavivirus infection elicits antibody responses against NS1. NS1-reactive antibodies and vaccination with NS1 have been shown to be protective in mouse models of flavivirus disease through Fc-dependent and Fc-independent mechanisms^10–15^. NS1 can act as a virulence factor that correlates with disease severity^16^ through several pathways, including activation of immune cells and platelets^9,17–21^. Importantly, NS1 can bind directly to endothelial cells. This leads to internalization and intracellular signaling with disruption of the endothelial glycocalyx layer (EGL) and intercellular junctions, ultimately resulting in endothelial barrier dysfunction and vascular leak^15,22–25^. Interestingly, NS1 from different flaviviruses can interact with endothelial cells and trigger endothelial barrier dysfunction with a tissue-specificity that reflects disease tropism; i.e., while DENV NS1 leads to dysfunction in lung endothelial cells (aligning with vascular dysfunction in the lung associated with severe dengue), WNV NS1 induces dysfunction specifically in brain endothelial cells (consistent with severe WNV encephalitis)^26^. We have previously identified molecular determinants on NS1 critical for pathogenesis through mutagenesis^10,27^, including the N-glycosylation mutant N207Q that is unable to trigger endothelial dysfunction^22^. Despite progress in recent years, the molecular mechanisms that underlie NS1-mediated endothelial barrier dysfunction and specifically the tissue tropism of NS1 and the functional impact of the N207Q mutant are currently not well understood.

In this study, we determined the interactome of DENV NS1 and compared it to the interactomes of WNV NS1 and DENV N207Q in human pulmonary endothelial cells to find functionally important host factors. We identified ephrin B1 (EFNB1), a transmembrane protein that can engage in bi-directional signaling, as a critical host factor in NS1-mediated endothelial barrier dysfunction and show that C-terminal phosphorylation of EFNB1 is required for NS1 to trigger hyperpermeability. Further, we demonstrated the direct binding of the EFNB1 receptor binding domain (RBD) to the wing and β- ladder domains of DENV NS1, but not WNV NS1, and show that the EFNB1 pathway can be targeted to prevent DENV NS1-induced vascular leak.

## Results

### DENV NS1 selectively interacts with host factors involved in intracellular signaling

We hypothesized that the NS1 N207Q mutant and WNV NS1 do not induce barrier dysfunction in human microvascular pulmonary endothelial cells (HPMECs) due to a failure to interact with critical host proteins. To identify functionally important host factors interacting with DENV NS1, we generated recombinant NS1 proteins that were C-terminally tagged with a hemagglutinin (HA) sequence (Fig. 1a, Fig S1a). In addition to the wildtype version of DENV NS1-HA (WT NS1^HA^), we included *Gaussia* luciferase HA (Gluc ^HA^) as an unrelated protein negative control in our analysis. To further delineate functionally important interaction partners, we included the mutant DENV NS1 N207Q-HA protein (N207Q NS1^HA^)^22^ and WNV NS1-HA (WNV NS1^HA^), which do not induce EGL disruption or hyperpermeability of HPMECs^26,27^, as non-phenotypic controls in our analysis (Fig. 1a). First, we tested whether the HA-tagged proteins retain their function. To do this, we assessed their capacity to disrupt the EGL through measurement of sialic acid and heparan sulfate on the cell surface and the induction of hyperpermeability through a reduction in transendothelial electrical resistance (TEER) across a monolayer of endothelial cells seeded in a Transwell system, as we have previously described for NS1^24^. We confirmed that the WT DENV NS1 and WT DENV NS1^HA^ triggered endothelial barrier dysfunction in HPMECs through the reduction of sialic acid levels on the cell surface and induction of hyperpermeability, while Gluc^HA^, DENV N207Q NS1^HA^ and WNV NS1^HA^ did not (Fig. 1b-d).

**Figure 1.**
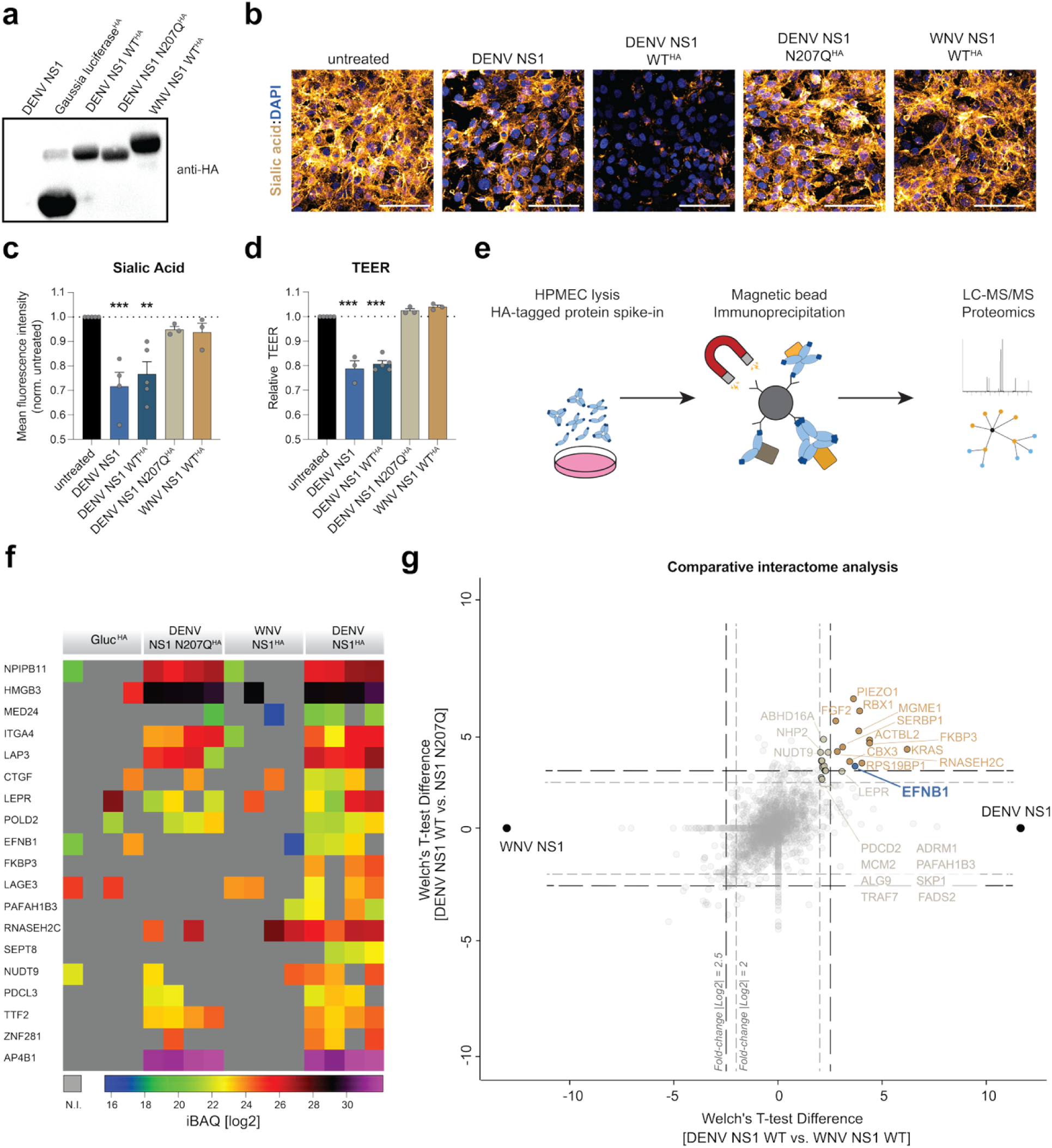
DENV NS1 selectively interacts with host factors involved in intracellular signaling. a, Western blot of Recombinant HA-tagged proteins using anti-HA antibodies. **b-c**, Sialic acid staining on the surface of HPMEC treated with the indicated recombinant proteins. Cells were treated, fixed and stained for sialic acid using wheat germ agglutinin (gold) and DAPI (blue). Representative micrographs are shown in b (scale bar, 100 μm) and fluorescence intensity quantification in **c** as the mean ± SEM (n=4). **d**, Endothelial permeability of a monolayer of treated with the indicated proteins. HPMEC were seeded in Transwells, treated and TEER measured at 0, 6 and 24 hours post treatment. Relative TEER at 6 hours is shown as mean ± SEM (n=4). **e**, Schematic of immunoprecipitation of HA- tagged NS1 from HPMEC lysates coupled to mass spectrometry (LC-MS/MS) bottom-up proteomics approach. **f**, Intensity-based absolute quantification (iBAQ) of selected host factors in each sample is illustrated as a heat map, with colors as indicated below the panel. Host proteins that were not identified (n.i.) in samples are shown in grey. **g**, Statistical analysis of host proteins in the DENV NS1 WT-treated sample compared to the non-phenotypic (DENV NS1 N207Q and WNV NS1) and negative (*Gaussia* luciferase, Gluc) controls. Host proteins with a fold-change of log2>2 are shown in taupe and those with an enrichment of log2>2.5 are shown in ochre. EFNB1 is indicated in blue. Statistical comparisons were performed by ordinary one-way ANOVA, with \*\**P* < 0.01, and \*\*\**P* < 0.001.

Next, we performed co-immunoprecipitation of HPMEC-derived host proteins with our HA-tagged baits and identified interaction partners through liquid-chromatography coupled to mass-spectrometry (LC-MS/MS) (Fig. 1e-h). Selection of proteins robustly enriched in DENV NS1 proteins when compared to Gluc or WNV NS1 controls identified 19 host proteins of interest (Fig. 1f). Analysis of gene ontology terms showed that these selected host proteins are involved in a range of biological processes but importantly included several factors that contain transmembrane domains and were reported to engage in intracellular signaling and cell adhesion (Fig. S1b). Notably, among these, we found a few host proteins selectively enriched in the WT NS1 sample compared to both our non-phenotypic controls (N207Q NS1 and WNV NS1) and therefore potentially involved in endothelial barrier dysfunction in HPMECs (Fig. 1g).

Among the cellular proteins distinctively enriched in DENV NS1, we identified ephrin B1 (EFNB1), a transmembrane protein expressed on the cell surface of endothelial cells from diverse tissues (Fig. 1g). EFNB1 is part of the ephrin family, which contains two distinct subgroups: the ephrin receptors (EphR) and their ephrin ligands (EFN). The latter group is further subdivided into the GPI-anchored ephrin A proteins and B-type ephrin ligands that contain a transmembrane domain and a C-terminal ‘post-synaptic density disc-large zo-1’ (PDZ)-binding domain. Interaction of EFNB1 with its cognate ephrin receptor (EphR) can lead to phosphorylation of EFNB1 on its C-terminus and engagement in bi-directional signaling. EFNB1 and its homologs EFNB2 and EFNB3 have previously been implicated in viral infections as receptors for henipaviruses, including Nipah, Hendra and Cedar viruses^28–31^. Altogether, we detected a range of host factors selectively interacting with DENV2 NS1 WT and focused our further analysis on EFNB1, a surface-expressed signaling molecule present in endothelial cells.

### Ephrin B1 is required for DENV NS1-mediated endothelial barrier dysfunction

We tested the functional requirement of EFNB1 through the generation of knockout (KO) HPMEC lines using a lentivirus-delivered CRISPR-Cas9 system with two guides targeting the *efnb1* gene and confirmed reduction in EFNB1 protein amounts by Western blot (Fig. 2a). EFNB1 KO cells were treated with NS1, and hyperpermeability was measured using the TEER assay. NS1 treatment resulted in a reduction in TEER in the non-targeting guide (NTG) control cells but did not affect barrier function when EFNB1 was depleted, indicating that EFNB1 plays a key role in NS1-mediated hyperpermeability (Fig. 2b). To ascertain specificity, we next tested if EFNB1 is generally involved in barrier function or if it is an NS1-specific host factor. We treated NTG and EFNB1 KO cells with two other activators of endothelial hyperpermeability, tumor necrosis factor-α (TNF-α) and Crimean Congo hemorrhagic fever virus glycoprotein 38 (GP38)^32,33^, and found comparable induction of hyperpermeability in the WT and EFNB1 KO cell lines, indicating that EFNB1 is specifically important for NS1-mediated endothelial barrier dysfunction.

**Figure 2.**
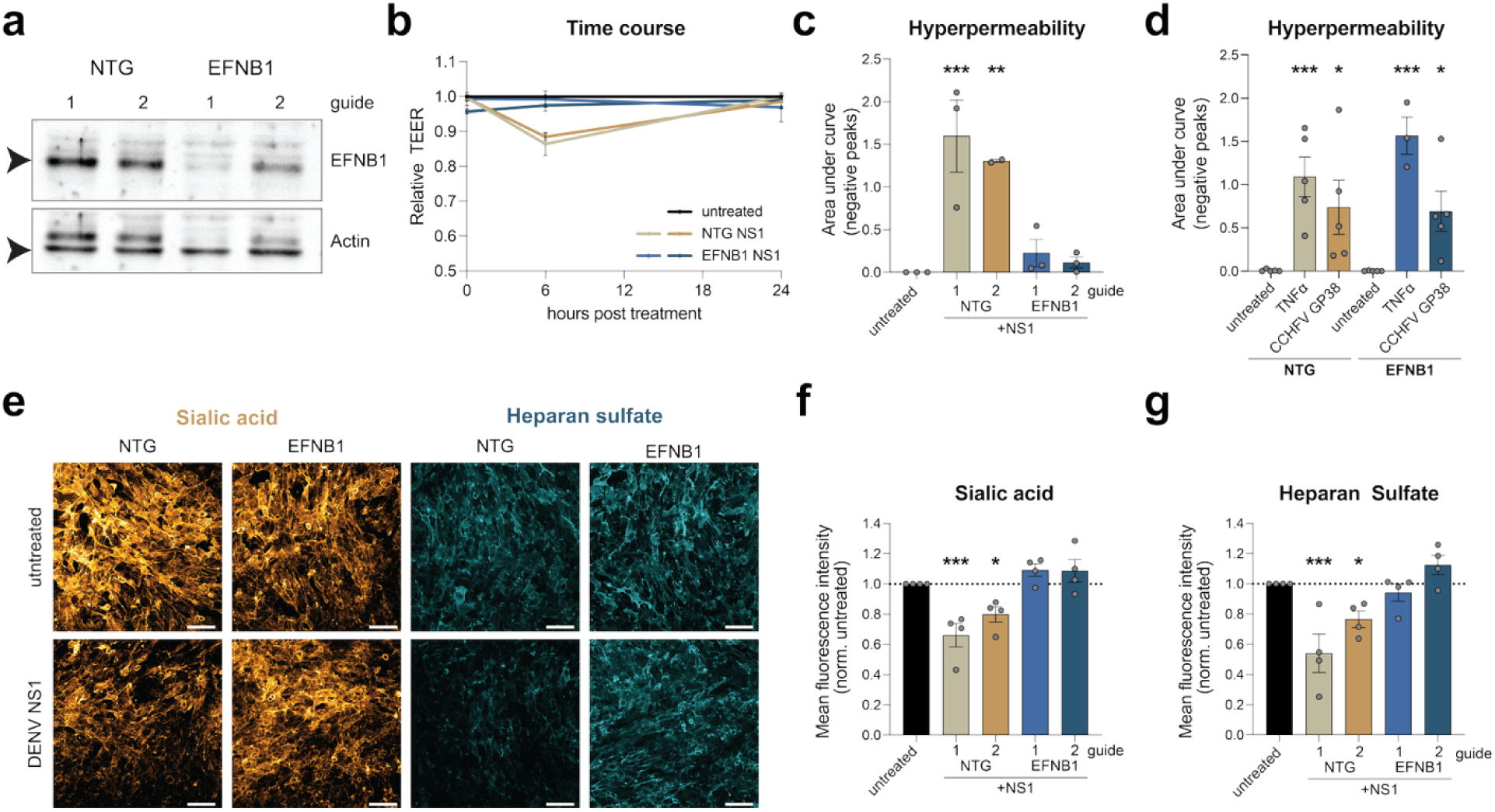
Ephrin B1 is required for DENV NS1-mediated endothelial barrier dysfunction. **a,** Western blot of EFNB1 knock-out (KO) cell lines with non-targeting guide (NTG) controls stained with mAbs against EFNB1 or β-actin. **b-c**, Endothelial permeability of NTG and EFNB1 KO cells treated with NS1. Relative TEER (**b**) and the area under the curve of the negative peaks (**c**) are shown as the mean ± SEM (n≥3). **d**, Endothelial permeability of NTG and EFNB1 KO cells treated with TNF-α and Crimean Congo hemorrhagic fever virus glycoprotein 38 (GP38). The area under the curve of the negative peaks is shown as the mean ± SEM (n≥3). **e-g**, Staining of EGL components of NTG control and EFNB1 treated with NS1. Cells were fixed and stained for sialic acid (gold, **e-f**) or heparan sulfate (cyan, **e** and **g**) for immunofluorescence assay (scale bar, 100 μm). Quantification of the mean fluorescence intensity is shown in **f** and **h** as the mean ± SEM (n≥3). Statistical comparisons were performed by ordinary one-way ANOVA, with \**P* < 0.05, \*\**P* < 0.01, and \*\*\**P* < 0.001.

Disruption of the EGL is a hallmark of endothelial barrier dysfunction; therefore, we treated NTG control and EFNB1 KO cells with NS1 and stained for sialic acid and heparan sulfate, which are major components of the EGL on the surface of HPMECs, as an orthogonal approach (Fig. 2e-g). NS1 treatment resulted in reduction in sialic acid and heparan sulfate staining in the NTG controls, as expected. In contrast, when EFNB1 was depleted, the EGL remained intact, indicating that EFNB1 is involved in the pathway leading to EGL disruption (Fig. 2e-g). In summary, these data demonstrate that EFNB1 is required for NS1-triggered endothelial hyperpermeability and EGL disruption.

### EFNB1 phosphorylation state regulates NS1-triggered endothelial barrier dysfunction

We next tested whether restoring protein expression of EFNB1 would recover the capacity of NS1 to induce endothelial barrier dysfunction in our HPMEC EFNB1 KO cells. To perform this experiment, we generated a lentivirus-delivered overexpression construct of EFNB1. EFNB1 is composed of an N-terminal extracellular receptor binding domain (RBD), a transmembrane domain, and a C-terminal signaling hub with a PDZ-binding domain (Fig. 3a). The C-terminus has been reported to be phosphorylated at six amino acid residues (Y313, Y317, Y324, Y329, Y343, Y344), and the two terminal phosphorylation sites of the PDZ-binding domain have been shown to be important for regulation of intracellular signaling pathways. Therefore, we introduced tyrosine (Y) to phenylalanine (F) substitutions in all tyrosine residues in the C-terminus to abrogate phosphorylation and test the role of this post-translational modification of EFNB1 in the NS1-mediated pathway. Further, we substituted the terminal tyrosine residues 343 and 344 with either phenylalanine to eliminate phosphorylation or with the negatively charged glutamic acid (E) to mimic constitutive phosphorylation. Finally, we added a C-terminal FLAG tag to allow for detection of EFNB1-FLAG expression using affinity tag-binding monoclonal antibodies (mAbs).

**Figure 3.**
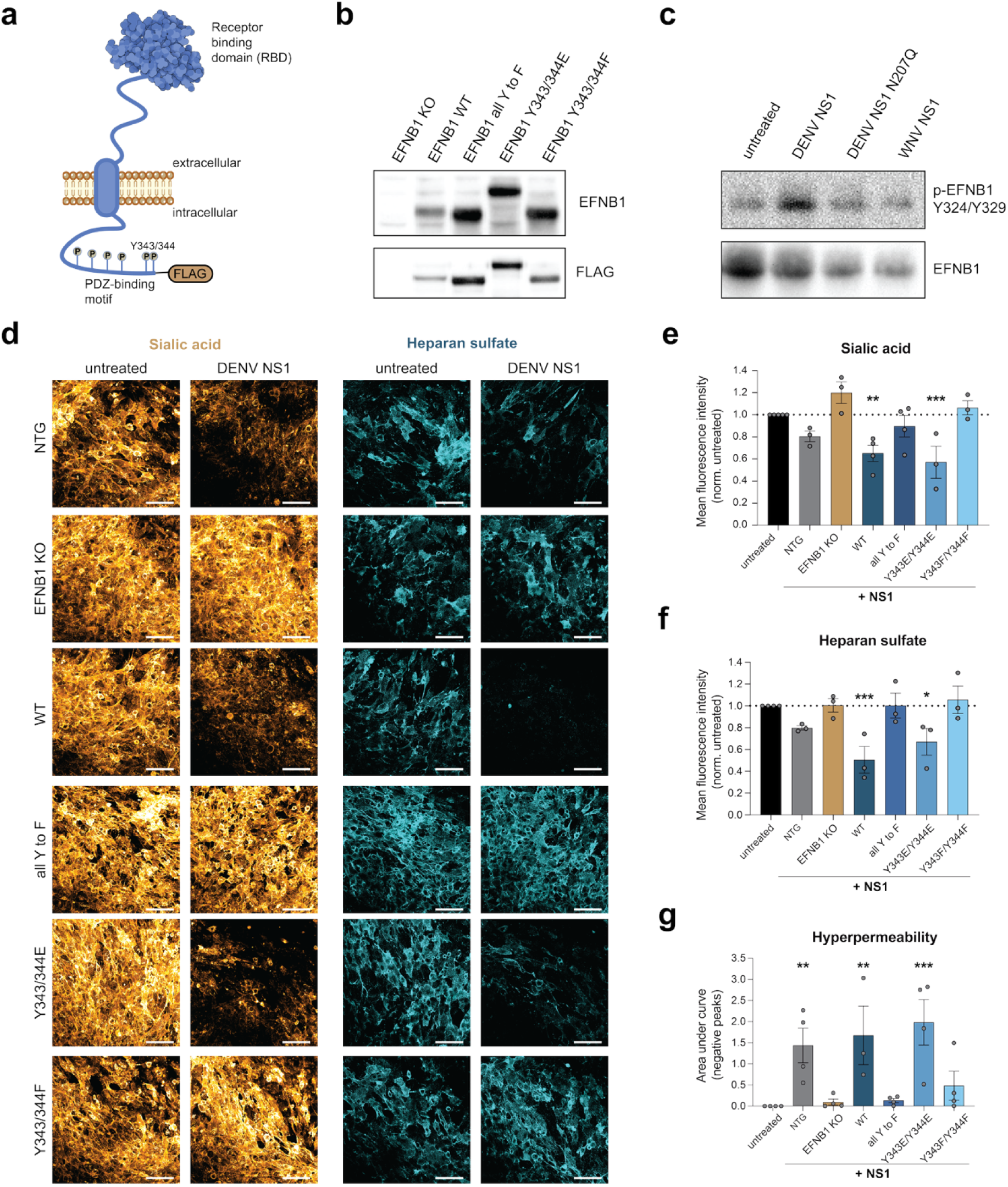
EFNB1 phosphorylation regulates NS1-triggered endothelial barrier dysfunction. **a,** Schematic of the EFNB1 protein at the plasma membrane showing the extracellular RBD and the intracellular PDZ-binding motif, with phosphorylation sites indicated by “P”. **b**, Western blot of lysates of EFNB1 KO cells either untreated or transduced with an expression construct encoding EFNB1 WT or mutants (all Y to F, Y343/344E, Y343/344F). Proteins were detected with an anti-EFNB1 antibody and an anti-FLAG-tag antibody. **c**, Western blot of samples from HPMECs treated with the indicated proteins. After treatment for 30 min, FLAG-tagged EFNB1 was isolated by immunoprecipitation. Proteins were detected using a mAb binding to EFNB1 phosphorylated at Y234/329 and an EFNB1- specific mAb. **d-f**, Staining of EGL components of EFNB1-reconstituted cell lines treated with NS1. Cells were fixed after 6 h and stained for sialic acid (gold) and heparan sulfate (cyan). Representative images are shown in **d** (scale bar, 100 μm) and the quantification of sialic acid and heparan sulfate staining is shown in **e** and **f** as the mean ± SEM (n≥3). **g**, Endothelial permeability of EFNB1- complemented cell lines treated with NS1. TEER was measured over a time course of 24 h and the area under the curve of the negative peaks was quantified as a proxy of hyperpermeability and shown as the mean ± SEM (n≥3). Statistical comparisons were performed by ordinary one-way ANOVA, with \**P* < 0.05, \*\**P* < 0.01, and \*\*\**P* < 0.001.

We first confirmed successful lentiviral transduction of the KO cell lines with the EFNB1 expression constructs by Western blot. While no protein was detectable in the EFNB1 KO cell line, we detected expression of an approximately 40 kDa protein by both the EFNB1 and FLAG antibodies for all the constructs, though the phosphor-mimetic EFNB1 migrated slightly slower (Fig. 3b). Next, we tested if NS1 treatment led to phosphorylation of EFNB1. We treated the EFNB1-FLAG-expressing HPMECs with DENV NS1 WT, DENV NS1 N207Q, or WNV NS1 or left the cells untreated and then lysed them 30 minutes post-treatment, since ephrin B phosphorylation post-stimulation was previously reported in this time frame^34^. Due to observed non-specific binding of the commercially available phospho- specific antibodies, we opted for purification of EFNB1 by anti-FLAG immunoprecipitation. After washing and elution, we used a mAb that binds to EFNB1 at phosphorylated residues Y324 and Y329. Upon DENV NS1 WT treatment, we observed an increase in phosphorylated EFNB1 compared to the untreated cells, which was not present in N207Q NS1- or WNV NS1-treated cells (Fig. 3c).

After observing phosphorylation of EFNB1 post-NS1 treatment, we tested whether reconstitution of EFNB1 expression in the EFNB1 KO cells by lentiviral transduction was able to restore EGL disruption by NS1 (Fig 3d-f). NS1 treatment led to reduction of both sialic acid and heparan sulfate on the surface of NTG-treated cells, whereas EFNB1 KO cells did not exhibit this reduction. Importantly, upon restoration of WT EFNB1 protein levels in the EFNB1 KO cells, NS1 did induce EGL disruption and, potentially due to higher expression levels, slightly increased the effect of NS1 treatment, confirming the functional importance of EFNB1 in the NS1 pathway. Further, expression of the exogenous phospho-dead EFNB1 could not restore the capacity of NS1 to induce EGL disruption, whereas NS1 treatment of the EFNB1 KO cell lines expressing the phospho-mimetic EFNB1 did show reduced sialic acid and heparan sulfate levels, indicating that mimicking constitutive phosphorylation of the C-terminal residues 343 and 344 restores NS1-mediated EGL disruption (Fig. 3d-f).

As an orthogonal approach, we performed TEER assays and ascertained that all the cell lines were able to form a comparable endothelial barrier. We observed overall similar TEER values at baseline across cell lines, although expression of the phospho-mimetic EFNB1 exhibited slightly lower barrier function at baseline (Fig. S2, p = 0.1896). Treatment of the NTG control cells with NS1 resulted in hyperpermeability, whereas the EFNB1 KO cells did not respond to NS1 treatment, as expected (Fig. 3g). Reconstitution with the EFNB1 WT construct recovered NS1-mediated endothelial dysfunction, while reconstitution with the phenylalanine substitution mutants of EFNB1 (all six and Y343/344F) did not. Interestingly, NS1 treatment of EFNB1 KO cells expressing the phospho-mimetic EFNB1 (Y343/344E) did result in hyperpermeability. Together, these data indicate that EFNB1 is a functionally important host factor and that C-terminal phosphorylation of EFNB1 regulates NS1- mediated endothelial barrier dysfunction.

### DENV NS1 directly binds to EFNB1

After identifying EFNB1 as a host factor that upon NS1 treatment is phosphorylated, we investigated whether NS1 engages in a direct protein-protein interaction with the EFNB1 RBD. We generated recombinant EFNB1 RBD proteins fused to the Fc region of a mouse IgG, an established tool in the field that improves EFNB1 stability and solubility, to test binding in a bead-based assay (Fig. 4a). NS1 from DENV2, WNV and ZIKV were coupled to beads, and the EFNB1 RBD or Gluc fused to the IgG Fc domain were added. We measured 40-fold higher binding of the EFNB1 RBD-Fc to DENV2 NS1 over the Gluc-Fc negative control (Fig. 4a-b). Consistent with the interactome data, we found that DENV2 NS1 exhibits significantly higher binding compared to WNV and ZIKV NS1 (Fig. 4a-b). Further, we measured comparable binding across the four DENV serotypes (Fig. 4c), with slightly higher binding observed for DENV1 and DENV3 (Fig. 4c). We also found that mutagenesis of the N-glycan site at position 207 resulted in reduced binding to EFNB1 RBD-Fc, in support of the interactome data.

**Figure 4:**
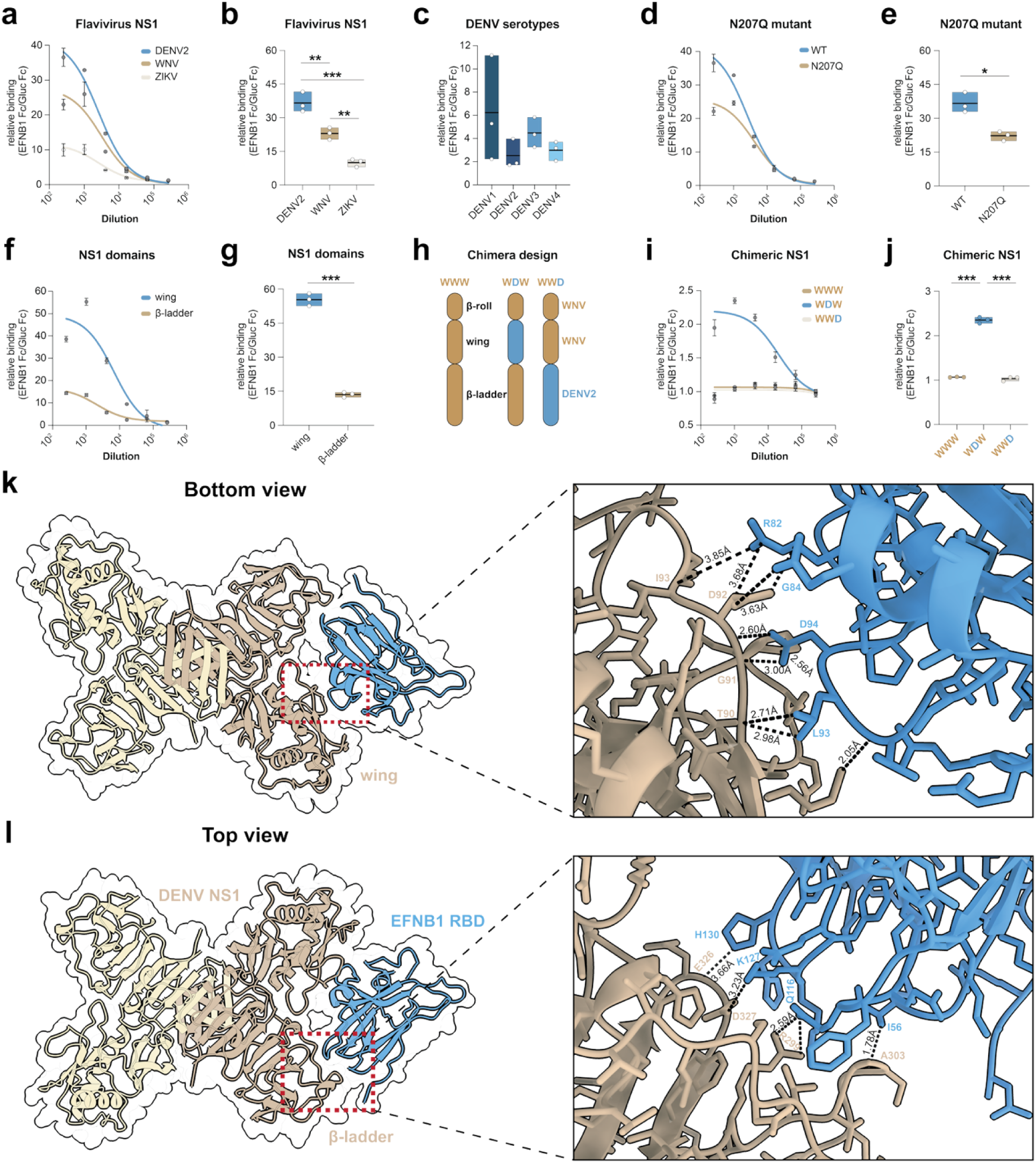
DENV NS1 wing and β-ladder domains bind to the EFNB1 RBD. **a-j,** Relative binding of EFNB1 Fc over Gluc Fc to flavivirus NS1 (**a-b**), DENV1-4 NS1 (**c**), DENV NS1 WT and N207Q (**d-e**), individual domains of DENV2 NS1 (**f-g**), or chimeric DENV-WNV NS1 (**i-j**). Luminex beads were coupled to the different antigens, and a dilution series of six 4-fold dilutions of the EFNB1 RBD Fc or the Gluc Fc fusion proteins was added. Binding was detected using fluorescently conjugated anti-mouse Fc antibodies, and the ratio of EFNB1 Fc MFI over the Gluc Fc MFI was plotted either as a curve with the mean ± SEM (**a, d, f, i**) or as a single point dilution at the peak of the curve with the mean as a bar (**b, c, e, g, j**) (n=3 technical replicates). Statistical comparisons were performed by t-test or ordinary one-way ANOVA, with \**P* < 0.05, \*\**P* < 0.01 and \*\*\**P* < 0.001. **h**, Schematic of the design of the chimeric NS1 proteins based on a WNV NS1 backbone (tan) with domain swaps of DENV2 NS1 (blue). **k-l**, Boltz2 structure prediction of the EFNB1 RBD (blue, UniProt: P98172) binding to DENV2 NS1 (tan, UniProt: P29990) as a cartoon representation. **k**, Dimeric NS1 shown from the bottom with an inset of the wing domain interacting with EFNB1. The major interacting residues and their distances are annotated. **l**, Dimeric NS1 shown from the top with an inset of the β-ladder domain interacting with the GH loop on EFNB1. The major interacting residues and their distances are annotated.

As an orthogonal approach to validate the Luminex bead-based approach, we inverted the orientation in an enzyme-linked immunosorbent assay (ELISA), coating the plate with the EFNB1 RBD-Fc fusion proteins and adding a dilution series of His-tagged Gluc, DENV2 NS1, WNV NS1, and ZIKV NS1 proteins. We detected no binding of Gluc to EFNB1 RBD Fc fusion and a significantly higher signal for DENV2 NS1 compared to WNV and ZIKV NS1 (Fig. S3a). Further, we interrogated the subcellular localization of EFNB1 and NS1 using the WT EFNB1-FLAG-expressing HPMECs. The subcellular distribution of EFNB1 and NS1 was in a punctate pattern with co-localization events where the peak intensity profiles of EFNB1 and DENV NS1 overlapped (Fig. 4f, Fig. S3b).

Next, we tested which domain of NS1 binds to the EFNB1 RBD utilizing two approaches. We first generated recombinant individual DENV NS1 wing and β-ladder domains and found that EFNB1, but not the Gluc Fc negative control, strongly bound to the wing domain, with lower but detectable binding to the β-ladder (Fig. 4f-g). Next, we generated domain swaps between DENV (D) and WNV (W), transplanting either the wing (WDW) or β-ladder domain (WWD) of DENV into the WNV backbone (Fig. 4h)^27^. Consistent with the previous results, while chimeras containing the DENV β-ladder (WWD) did not increase binding compared to wildtype WNV NS1, when the wing domain was introduced into a WNV backbone (WDW), we observed significantly increased binding of EFNB1 RBD compared to the negative control (Fig. 4i-j). Together with the individual domains, these data indicate that the wing domain is the main driver of the EFNB1-NS1 interaction, with a minor contribution by the β-ladder domain.

Next, we used the structure prediction model Boltz2^35^ to investigate the potential structure of the NS1-EFNB1 complex (Fig. 4k-l, Fig. S3c-d). While Boltz2 predicted several different conformations, the highest scoring structural prediction (confidence score 0.87, interface Predicted Template Modeling = 0.79) showed the EFNB1 RBD (UniProt: P98172) binding to the surface between the wing and β-ladder domains of DENV2 NS1 (UniProt: P29990), a site that is accessible in all multimeric states^6,7^. The major residues predicted to drive the interaction are located in a loop of the EFNB1 RBD (residues R82, G84, D94 and L93) binding to a loop in the wing domain of NS1 (residues T90, G91, D92 and I93) (Fig. 4k). The minor interaction of the EFNB1 RBD binding to the β-ladder domain is driven by residues in the GH loop of EFNB1 (Q116, K127, H130) binding to residues R299, A303 as well as E326 and D327 on NS1 (Fig. 4l). To summarize, by employing several orthogonal biochemical and computational modelling approaches, we showed that EFNB1 directly binds to the wing and β- ladder domains of NS1.

### EFNB1 can be targeted to prevent endothelial barrier dysfunction

The direct interaction of EFNB1 and NS1 led us to investigate whether EFNB1 can represent a target to block vascular dysfunction either using small molecule inhibitors or using the EFNB1 Fc fusion protein as a decoy to prevent endothelial cell binding (Fig. 5a). EFNB1 phosphorylation can lead to signaling through a myriad of pathways, including protein kinase C (PKC) signaling^36–38^. We used the pan-PKC inhibitor Go 6983^39^ and observed a dose-dependent inhibition of NS1-triggered endothelial hyperpermeability, with an IC_50_ of approximately 10 nM (2 to 42 nM) (Fig. 5b).

**Figure 5:**
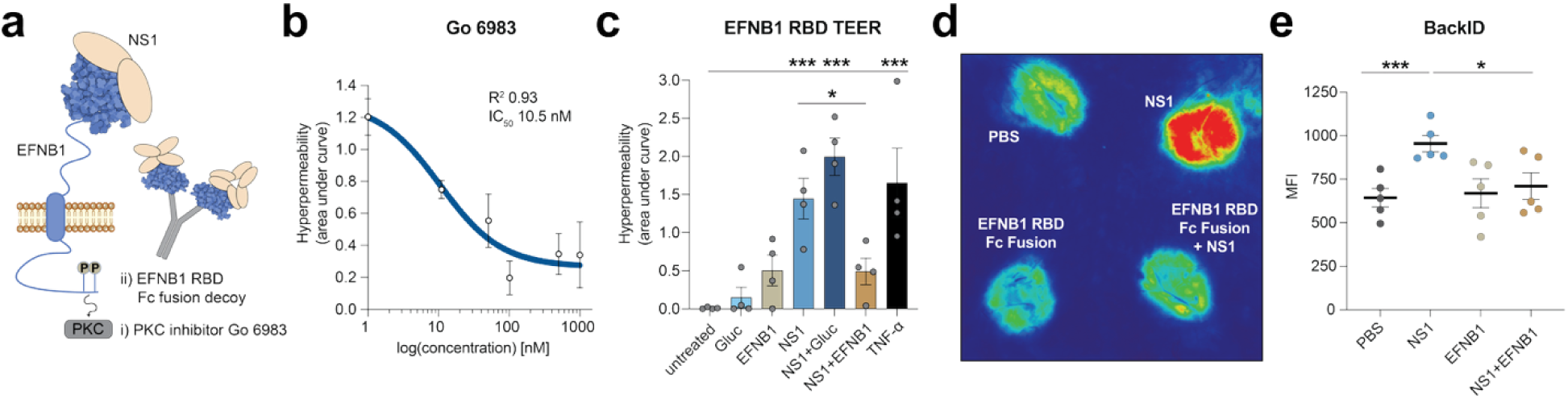
EFNB1 binds to NS1 and can be targeted to prevent endothelial dysfunction. **a,** Schematic of the tested approaches to target EFNB1. **b**, Endothelial permeability of HPMEC treated with NS1 in presence of decreasing concentrations of the PKC inhibitor Go 6983. The area under the curve of the negative peaks was calculated and plotted against the inhibitor concentration in nM as the mean ± SEM (n≥2). **c**, Endothelial permeability of HPMEC treated with NS1 or with a combination of NS1 and Fc fusion proteins. NS1 was complexed with EFNB1 RBD Fc fusion proteins or as a negative control with the Gluc Fc fusion protein and TEER was measured. The area under the curve of the negative peaks was quantified as the mean ± SEM (n≥3). **d-e**, Vascular leak in the mouse dermis. NS1 was complexed with the EFNB1 RBD Fc fusion proteins and the complex or the individual proteins were injected intradermally into the shaved skin of C57BL/6 mice. PBS was used as a negative control. The tracer dye Dextran-680 was simultaneously injected intravenously, and dye extravasation was visualized using a LiCor Scanner after extraction of the skin. A representative scan is shown in **d** and the quantification of the mean fluorescent intensity (MFI) of the extravasated tracer dye is shown in **e** as the mean ± SEM (n=5). Statistical comparisons were performed by ordinary one-way ANOVA, with \**P* < 0.05, \*\**P* < 0.01, and \*\*\**P* < 0.001.

Next, NS1 was complexed either with the EFNB1 RBD-Fc or with the Gluc-Fc protein, and TEER assays were conducted (Fig. 5c). Whereas untreated cells and cells treated with the Fc fusion proteins alone did not exhibit hyperpermeability, NS1 and TNF-α triggered a reduction in barrier function (Fig. 5c). Notably, NS1 induced endothelial hyperpermeability when co-incubated with the Gluc-Fc negative control, but hyperpermeability was prevented when the NS1-EFNB1-Fc complexes were added to the cells, indicating that EFNB1 RBD-Fc fusion protein can act as a decoy to block NS1-mediated endothelial dysfunction. Finally, we tested if the EFNB1-Fc decoy could inhibit vascular leak in our localized murine dermal leak model. Mice were intradermally injected with DENV NS1, the EFNB1-Fc fusion protein, or the combination of NS1 and EFNB1 RBD-Fc, and extravasation of the intravenously injected tracer dye Dextran-680 was measured as a proxy for vascular leak. As expected, NS1 triggered an increase in Dextran-680 extravasation compared to PBS injection, while the NS1-EFNB1-Fc fusion protein alone did not (Fig. 5d-e). Moreover, the EFNB1-Fc proteins in complex with NS1 prevented vascular leak induction, confirming our *in vitro* results. In summary, we find that EFNB1 can be targeted by small molecule inhibitors or Fc fusion decoys to prevent endothelial dysfunction and vascular leak.

## Discussion

In this study, we identified EFNB1 as a surface-expressed binding partner involved in DENV NS1- mediated endothelial barrier dysfunction and vascular leak. Using a comparative proteomic approach, we found that DENV NS1 specifically interacts with several host factors engaging in intracellular signaling, including the transmembrane protein EFNB1. Genetic and biochemical assays revealed that NS1-mediated endothelial dysfunction is dependent on C-terminal phosphorylation of EFNB1. Further, we showed that the EFNB1 RBD binds directly to DENV NS1 and that EFNB1 can be targeted to prevent vascular leak.

Previous studies have interrogated potential host interaction partners of flavivirus NS1 but generally relied on overexpression of NS1 in kidney or liver cells and identified mostly intracellular interaction partners^40–42^. We were specifically interested in host factors involved in NS1-mediated endothelial barrier dysfunction, which is why we opted to conduct our interactome analysis with HPMEC. The comparison with the two non-phenotypic controls, a closely related flavivirus NS1 and a point mutant of DENV NS1 that do not trigger the NS1-mediated pathogenesis cascade, allowed for the stringent selection of potentially functionally important host factors. Our identification of signaling molecules such as EFNB1 is consistent with previous studies that demonstrated that NS1 is internalized and triggers intracellular signaling^9,17,22,24,43^, although there are no previous reports of EFNB1 as a potential DENV NS1-interacting host factor. Besides EFNB1, several other host proteins of interest were identified as selective WT DENV NS1 interactors, including the leptin receptor (LEPR), integrin alpha 4 (ITGA4), and phosducin-like protein 3 (PDCL3), which are all involved in intracellular signaling. Their potential contribution to NS1 pathogenesis will be further characterized in future studies.

EFNB1 is a single-pass membrane protein that has been shown to be involved in a variety of biological processes – importantly, including intracellular signaling, modulation of tight and adherens junctions, and as a receptor for Cedar virus^29,37,38,44^. In addition to EFNB1, humans express the homologs EFNB2 and EFNB3; interestingly, the EFNB homologs are expressed in endothelial cells with a distinct tissue- specific expression pattern, where EFNB1 and EFNB2 exhibit broad expression while EFNB3 expression is mostly limited to the brain. In agreement with the reported expression patterns, we only detected EFNB1 and EFNB2 in the lung endothelial cell lysates, and only EFNB1 was detected in the IP samples, indicating that EFNB2 might not interact with DENV NS1. Additionally, the expression pattern of the EFNB homologs is consistent with the tropism of flavivirus NS1, i.e., DENV NS1 has broad tissue tropism and induces dysfunction in endothelial cells from multiple organs, while WNV NS1 only causes dysfunction in brain-derived endothelial cells^26^. It will be important to investigate whether EFNB1-3 expression and activation can contribute to the observed tropism of flavivirus NS1 pathogenesis, as might be suggested by the predicted interaction site of EFNB1 on NS1 that had previously been shown to determine endothelial cell tropism^27^.

In this study, we show that EFNB1 binding to NS1 triggers NS1-mediated endothelial dysfunction and that direct protein-protein interaction leads to intracellular signaling. The structural prediction of the NS1-EFNB1 complex indicated that both the wing and the β-ladder domain on NS1 contribute to the interaction, which is consistent with the biochemical data that showed approximately 50- and 15-fold higher relative binding of EFNB1 RBD to the wing and the β-ladder domains, respectively. The main drivers of the interaction in the wing domain are a stretch of four amino acids (T90, G91, D92, I93), which are highly divergent in WNV (V90, E91, K92, Q93). Intriguingly, a previous study from our group identified the GDI motif to be the molecular determinants of WNV versus DENV NS1 tropism^27^, indicating that EFNB1 might be a key host determinant of the observed tissue tropism of flavivirus NS1. The predicted interacting residues on the β-ladder domain (R299, A303, E326, K327) are key residues in the epitope of the protective mAb 2B7, indicating that the mechanism of action of 2B7 might be steric hindrance of the EFNB1 binding site. Interestingly, 2B7 has been shown to bind promiscuously to many flavivirus NS1 proteins, and R299 is a highly conserved residue across the flavivirus genus^10^ – which might explain the observed reduced but detectable binding of WNV and ZIKV (Fig. 4a-b) despite their sequence differences in the GDI loop on the wing. Interestingly, the remaining interacting residues are the most variable residues in the 2B7 epitope, and when these residues were mutated in our previous study, NS1 retained its capacity to bind to HPMEC, likely via the wing domain, but exhibited a reduction in hyperpermeability induction^10^. This suggests a model where the binding of both the wing and β-ladder domains to EFNB1 is needed for the full activation of the pathway and supports the central role of EFNB1 in NS1-mediated endothelial barrier dysfunction.

Previous studies have investigated host factors potentially involved in NS1 attachment and pathogenesis. For example, toll like receptor 4 (TLR4) was shown to be involved in NS1-mediated pro- inflammatory cytokine production and thrombocytopenia^18,45^, although subsequent studies did not find differences in vascular leak upon NS1 treatment in TLR4-deficient mice^33^. Additionally, a recent study indicated scavenger receptor B1 (SRB1) as a potential interaction partner^46^. It is currently unclear how these different host proteins contribute to the overall pathogenesis and whether multiple proteins can bind to NS1 to mediate endothelial dysfunction. Validation of the relative contribution of these host proteins in genetically deficient animals is a key next step. In the case of EFNB1, testing in EFNB1-deficient animals is challenging due to its essential role in development^47^, although we were able to demonstrate that an EFNB1 RBD decoy can prevent vascular leak *in vivo*.

We found that upon NS1 treatment, EFNB1 was C-terminally phosphorylated, which was required for NS1-mediated endothelial barrier dysfunction. Previous studies of EFNB1 signaling provide evidence that EFNB1 phosphorylation can lead to both disruption of cell adhesion and reorganization of tight junctions^36,44,48^. It is currently unclear whether NS1 treatment leads to a change in the interaction partners of EFNB1 and which signaling proteins might be recruited or dispersed after NS1 interaction. We assessed the potential of EFNB1 as an intervention target using a small molecule inhibitor of downstream pathways and by blockade with a soluble EFNB1 RBD acting as a decoy to prevent vascular leak *in vitro* and *in vivo*. Notably, analysis of differentially regulated genes during the acute phase of DENV infection in human PBMCs has revealed a significant upregulation of the ephrin receptor B2 (EPHB2), of which EFNB1 is a ligand, indicating that the ephrin pathway is dysregulated in acute dengue^49–51^. Interestingly, the ephrin/EPH family of proteins has been previously implicated in vascular leak during sepsis, and treatment with an EphA4 Fc fusion protein alleviated vascular leak in mouse models of sepsis^52–54^, indicating that this family of proteins are regulators of endothelial barrier function and could represent an attractive target for therapeutic intervention.

Limitations of this study include that EFNB1 might not be the only surface-expressed host protein that is required for NS1-mediated endothelial dysfunction, and future studies, especially in *in vivo* models, are needed to evaluate the relative contribution of candidate proteins. Further, while structural predictions have been shown to be reliable, especially when interactions are validated through experimental data, experimentally resolving the structure of the EFNB1-NS1 complex will be important and could be critical for vaccine and mAb design.

In summary, we identified EFNB1 as critical host factor in NS1-induced endothelial barrier dysfunction. This study builds a model where circulating NS1 binds EFNB1 on endothelial cells, leading to phosphorylation, ultimately resulting in endothelial barrier dysfunction. The data provided here are important for vaccine and antigen design as well as the development of interventions targeting the ephrin family for preventing NS1-mediated vascular leak, a hallmark of severe DENV infection, as demonstrated through our inhibitor and EFNB1 RBD decoy approaches. Dengue is one of the largest global public health threats, and this study can contribute to the ongoing efforts for new and better preventative and therapeutic interventions.

## Materials and Methods

### Ethics statement

Experiments conducted at UC Berkeley were pre-approved by the UC Berkeley Animal Care and Use Committee (Protocol AUP-2014-08-6638-3) and conducted in compliance with Federal and University regulations. Six-week-old wild-type C57BL/6J female mice were purchased from Jackson Laboratory (strain 000664) and housed under specific pathogen-free conditions at the UC Berkeley Animal Facility.

### Cell culture

Human pulmonary microvascular endothelial cells (HPMEC) [line HpMEC-ST1.6 R] were a gift from Dr. J.C. Kirkpatrick at Johannes Gutenberg University, Germany^55^. HPMEC were cultured in endothelial growth basal medium 2 (EGM-2) with an EGM-2 microvascular cell supplemental bullet kit (Lonza) and maintained at 37°C with 5% CO_2_. HEK293T cells were cultured in Dulbecco’s modified Eagle medium (DMEM, Gibco) with 5% fetal bovine serum (FBS; Gibco) and maintained at 37°C with 5% CO_2_. Expi293F cells were cultured in Expi293™ Expression Medium (Gibco) and maintained at 37°C with 8% CO_2_.

### Cloning

The Gluc^HA^, and Fc fusion protein sequences were codon-optimized and synthesized by Twist Biosciences. The HA tag was added to the C-terminus of existing DENV and WNV NS1 plasmid sequences which already contained a C-terminal 6x histidine tag for purification using PCR with overlap primers (DENV: tttctcgagtcaagcgtaatctggaacatcgtatgggtaGTGATGGTGATGGTGATGagctgtg, WNV: tttctcgagtcaagcgtaatctggaacatcgtatgggtagtgatggtgatggtgatgAGCATTC). HPMEC EFNB1 KO cells were complemented with three EFNB1 mutant constructs, in addition to wildtype EFNB1 as a positive control. A phosphorylation mimic (Y343E, Y344E) was created by substituting the two PDZ tyrosine phosphorylation sites with a glutamate residue. A phosphorylation dead mutant abolished the PDZ domain phosphorylation sites by substituting tyrosine for phenylalanine (Y343F, Y344F). Lastly, a variant with all phosphorylation sites removed was created by mutating all 6 tyrosine phosphorylation sites to phenylalanine (Y313F, Y317F, Y324F, Y329F, Y343F, Y344F). The sequences encoding these EFNB1 mutants were synthesized by Twist Bioscience and transferred to the pENTR vector by restriction cloning before transfer to the pLX301 lentiviral vector^56^ via Golden Gate cloning to create plasmids containing the mutant EFNB1 sequence with a CMV promoter as well as a C-terminal FLAG tag.

To generate Fc-fusion proteins, we cloned the receptor-binding domains (RBDs) of Ephrin-B1 and of the Gluc sequence into expression vectors containing the Fc domain of murine IgG2b. EFNB1–3 RBD fragments and vector backbones were digested with appropriate restriction enzymes (EcoRI and NheI) and ligated to generate the final fusion constructs. Cloning success was verified by Sanger sequencing.

### Lentivirus production and transduction

Lentiviruses encoding the mutant EFNB1 variants were produced as previously described^57^. HEK293T cells were seeded at 1,000,000 cells per well and transfected the next day with 2 μg of either the pLentiCRISPR^58^ or pLX301 plasmids containing the guides (EFNB1: guide1: caccgACGTGTTGGTCACCTGCAAT/aaacATTGCAGGTGACCAACACGTc; guide 2: caccgGAGTTCAGCCCCAACTACAT/aaacATGTAGTTGGGGCTGAACTCc) or the sequence of the gene of interest together with 1.3 μg and 0.7 μg of the psPAX2 and PMD2.G packaging vectors, respectively. Supernatants were collected every 8 hours, filtered, and used for reverse transduction of endothelial cells. Transduced cells were allowed to recover and then were selected using puromycin (20 μg/mL). Expression levels after knockout or overexpression were verified using Western blot analysis with specific antibodies.

### Recombinant protein production

Recombinant proteins were either purchased from Native Antigen or produced in-house. Chimeric DENV2-WNV NS1 proteins were constructed as described previously and labelled as WDW for the substitution of the WNV wing domain with the DENV2 wing domain (residues 37-152) or WWD for the substitution of the WNV β-ladder domain with the DENV2 β-ladder domain (residues 180-352)^27^. Recombinant DENV2 NS1 wing-Halotag (residues 30-174) and DENV2 NS1 β-ladder-Halotag domains (residues 178-352) contain a 3x(GGGGS) linker between the NS1 domain and the Halotag and an 8x- histidine tag on the C terminal. All in-house produced proteins were generated by transfection of Expi293F™ cells using ExpiFectamine™ 293, following the manufacturer’s protocol. Briefly, cells were seeded at 3,000,000 cells/mL and transfected with 1 µg of plasmid DNA per mL of culture. DNA–lipid complexes were formed in Opti-MEM™ and added to the cells, followed by the addition of Enhancers 1 and 2 the next day. Supernatants were harvested 5–6 days post-transfection. The 6X-histidine- and HA-tagged proteins were purified using Cobalt beads (25229, ThermoFisher Scientific). Transfected cell supernatants were mixed with beads in washing buffer (20 mM sodium phosphate, 500 mM sodium chloride, 20 mM imidazole, pH 7.4) and incubated overnight shaking at 4°C. Beads were loaded on a gravity flow column and washed with three column volumes of washing buffer before addition of 5 mL of elution buffer (20 mM sodium phosphate, 500 mM sodium chloride, 300 mM imidazole; pH 7.4) and collection in 1-mL fractions. Fractions were analyzed by Coomassie blue staining to evaluate protein content, and recombinant protein containing fractions were pooled and concentrated using Amicon filtration (10 kDa). Fc-fusion proteins were purified from culture supernatants using Protein G affinity chromatography on an Akta FPLC system (Cytiva). EFNB1-Fc fusion proteins typically eluted as a sharp peak around 390 mL. Pooled elution fractions from all recombinant proteins were further purified by size exclusion chromatography to isolate monomeric proteins needed for Luminex bead coupling. Protein concentration was measured by absorbance at 280 nm and Pierce™ Bicinchoninic Acid Protein Assay Kits (23225, ThermoFisher Scientific).

### Immunoprecipitation and Western blot

All immunoprecipitations were performed on ice and in the presence of protease inhibitors (cOmplete EDTA-free protease inhibitor cocktail, Sigma-Aldrich) under conditions described previously^59^. For the comparative interactome analysis, HPMEC were seeded at 2,000,000 per 15-cm dishes, grown until confluence, and washed with PBS before scraping in 5 mL of PBS. Cells were centrifuged, and PBS was removed before freezing of cell pellets at -80°C. Three cell pellets were combined and resuspended in 400 μL of lysis buffer (50 mM Tris-HCl, 150 mM NaCl, 1 mM EDTA, 0.5% dodecylmaltoside, 5% glycerol). Lysis was performed for 30 minutes (min) on ice with regular inverting before centrifugation at 15,000 x g for 15 min at 4°C. Supernatants were transferred to new tubes and mixed with 15 µg of the HA-tagged recombinant proteins. Complexes were allowed to form overnight at 4°C under rotation before addition of 100 µL Pierce™ Anti-HA Magnetic Beads (ThermoFisher Scientific) and further incubation for three hours under rotation at 4°C. Beads were separated using a magnet and washed four times using the lysis buffer and three additional times using PBS. Elution buffer (200 μL, 3% SDS in 50 mM Tris-HCl pH 7.4) was added and incubated for 5 min.

For the analysis of the phosphorylation state of EFNB1, HPMEC WT EFNB1-FLAG-expressing cells were grown until confluent in 6-well tissue-culture plates and treated with 10μg/mL of DENV NS1, WNV NS1, or DENV N207Q NS1, or left untreated as a negative control. After 30 min, cells were washed twice with TBS (DEFINE) and lysed with radioimmunoprecipitation buffer (ThermoFisher Scientific) containing a phosphatase inhibitor cocktail (Cell Signaling). The plate was incubated with shaking at 4°C for 20 min before centrifugation at 16,000 x g for 10 min at 4°C, followed by collection of the supernatant. Immunoprecipitation of EFNB1-FLAG was performed by incubating 5 μg of total protein per sample with anti-FLAG beads at 4°C overnight, followed by three washes with TBS. Proteins were then eluted by boiling for 10 minutes at 95°C in 2X Laemmeli sample buffer (0.1 M Tris [pH 6.8], 4% SDS, 4 mM EDTA, 0.286 M 2-mercaptoethanol, 10 mL of 3.2 M glycerol, 0.05% bromophenol blue). The boiled samples were separated on a 12% gradient SDS-PAGE. Immunoblotting was performed using a primary antibody specific for EFNB1 tyrosine phosphorylation at residues 324 and 329 (1:1000; Cell Signaling) or an anti-EFNB1 antibody (1:1,000; Cell Signaling). Primary antibodies were incubated at 4°C overnight and then incubated with a species-specific anti-IgG secondary antibody conjugated to horseradish peroxidase. Protein visualization was performed using the ChemiDoc Imager (Bio-Rad) after washing three times with 0.1% Tween20 in TBS (TBS-T).

### Sample preparation for quantitative LC-MS/MS proteomics

To characterize the interactome of DENV NS1-WT-HA, DENV NS1-N207Q-HA, WNV NS1-WT-HA and Gaussia-HA, anti-HA immunoprecipitation was performed using endothelial HPMEC cell lysates as described above. Whole cell lysate (50 µg) and eluate samples were processed for quantitative LC-MS/MS proteomics (n=4). The samples were precipitated twice with acetone and resuspended and denatured in 40 µL U/T buffer (6 M Urea/2 M thiourea in 10 mM HEPES, pH 8.0). Proteins were reduced and alkylated in 10 mM DTT and 55 mM iodoacetamide, followed by digestion with 0.5 µg (eluates) or 1 µg (inputs; 50 µg protein) of LysC (FUJIFILM Wako Chemicals) and trypsin (Promega), respectively, in ABC buffer (50 mM NH_4_HCO_3_ in water, pH 8.0) overnight at 25°C, shaking at 800 rpm. After digestion, peptides were purified on stage tips with 3 layers of C18 Empore filter discs (3M) as previously described^60^.

Samples were analyzed on a Vanquish Neo LC system (ThermoFisher Scientific) coupled to a Orbitrap Exploris 480 mass spectrometer (ThermoFisher Scientific) equipped with a Nanospray Flex source (ThermoFisher Scientific). Peptides were injected into an Acclaim PepMap 100 trap column (2 cm × 75 μm, 3 μm C18; ThermoFisher Scientific) and next separated on a 25 cm × 75 μm column (1.7 µm C18 beads UHPLC column; Aurora Ultimate) with a packed emitter tip (Ion Opticks), at a constant flow rate of 300nl/min over a 100-min linear gradient. The column temperature was maintained at 50°C using an integrated column oven (Sonation GmbH). The column was equilibrated using 3 column volumes before loading samples in 100% buffer A (99.9% Milli-Q water, 0.1% formic acid (FA)). Samples were separated using a linear gradient from 9 to 31% buffer B (99.9% ACN, 0.1% FA) over 56 min before ramping up to 43% (35 min), 100% (1 min), and then sustained for 8 min. The Orbitrap Exploris 480 was operated in positive ion mode, with a positive ion voltage of 2000 V in data-dependent acquisition mode (DDA) using the Thermo Xcalibur software (v. 4.5.474.0). DDA analysis was performed with a cycle time of 1.5 seconds. Survey scans were acquired at 120,000 resolution, with a full scan range of 350-1400 m/z, an automatic gain control (AGC) target of 300%, and a maximum ion injection time of 25 ms, intensity threshold of 5 x 10^3^, 2–6 charge state, dynamic exclusion of 90 seconds, and mass tolerance of 10 ppm. The selected precursor ions were isolated in a window of 1.6 m/z, fragmented by a higher-energy collisional dissociation (HCD) of 30. Fragment scans were performed at 15,000 resolution, with an Xcalibur-automated maximum injection time and standard AGC target.

### Raw mass spectrometry data processing and analysis

Raw MS data were processed with the MaxQuant software v. 2.2.0.0 using the built-in label-free quantitation algorithm and Andromeda search engine (PMID: 27809316). The search was done against the *Homo sapiens* proteome (UniprotKB release UP000005640; Taxon ID 9606) containing forward and reverse sequences plus the sequences of *Gaussia*-His-HA (PDB: 7D2O), WNV NS1-His-HA (WNV-1/US/BID-V4702/2002), DENV2 NS1-His-HA (DENV2_16681), and DENV2 NS1-N207Q-His-HA. Additionally, the intensity-based absolute quantification (iBAQ) algorithm and match between runs option were used. In MaxQuant, carbamidomethylation was set as fixed and methionine oxidation and N-acetylation as variable modifications. Initial search peptide tolerance was set at 20 p.p.m., and the main search was set at 4.5 p.p.m.. Experiment type was set as data-dependent acquisition with no modification to the default settings. Search results were filtered with a false discovery rate of 0.01 for peptide and protein identification. The Perseus software v.1.6.15.0 was used to further process the affinity-purification datasets. Protein tables were filtered to eliminate the identifications from the reverse database and common contaminants. In the subsequent MS data analysis, only proteins identified on the basis of at least one peptide and a minimum of three quantitation events in at least one experimental group were considered. The iBAQ protein intensity values of the interactome dataset were median-normalized, log_2_-transformed, and missing values were filled by imputation with random numbers drawn from a normal distribution calculated for each sample (PMID: 27348712). DENV NS1 selective interactors were determined by two-sided Welch’s T-tests with permutation-based false discovery rate (FDR) statistics (250 permutations, FDR threshold 0.05) or using a log_2_(fold-change) ≥ 2 or 2.5 cut-off as indicated in the corresponding figures. Individual profiles of normalized intensities within unimputed matrix were visually assessed to further shortlist candidate proteins. Only proteins consistently identified in at least 3/4 biological replicates of either DENV NS1 WT or NS1-N207Q and displaying a log_2_(fold-change) ≥ 10 over negative controls (Gluc or NS1 WNV) were retained for further analysis. Results were plotted as scatter plots and heat maps using Perseus (PMID: 27348712).

### Transendothelial electrical resistance (TEER) assay

To measure endothelial permeability, 50,000 cells were seeded in the apical chamber of a 24-well Transwell plate (Transwell permeable support, 0.4 μM, 6.5 mm insert; Corning). Next, 1.5mL of EGM- 2 medium was added to the lower chamber, and 300 uL was added to the apical chamber. Half the volume of medium was changed in both chambers daily for 3 days until the cells reached a confluent monolayer. Electrical resistance was measured immediately before treatment, at 6 hours post-treatment, and at 24 hours post-treatment. Unless indicated otherwise, NS1 was used at 6 μg/mL, CCHFV GP38 at 2.5 μg/mL, and TNFα at 10 ng/mL. STX4 electrodes were used to measure the resistance in ohms (Ω) between the apical and basal compartments using an Epithelial Volt Ohm Meter (EVOM) Manual (World Precision Instruments). Medium-only wells were used to calculate the baseline electrical resistance, and TEER measurements were normalized to the electrical resistance of untreated cells. The area under the curve of the negative peaks was calculated using the t=0 and t=24 time-points for each well as the baseline. To determine the IC_50_ of GO 6983, the inhibitor was diluted in PBS, and 10 nM, 50 nM, 100 nM or 1000 nM were added to the upper chamber in addition to 6 μg/mL of DENV NS1. The area under the curve of the negative peaks was calculated as above, and a three-parameter nonlinear regression model was used to fit a curve with the AUC values of the negative control set as the bottom constant (R^2^ = 0.9285).

### Immunofluorescence assay

To measure EGL components on the surface of endothelial cells, 50,000 cells were seeded per well on glass coverslips in a 24-well plate. Medium was exchanged daily for 2-3 days until cells were confluent for 24 hours. Then, medium was exchanged, and cells were treated with 6 ug/mL of recombinant proteins. Cells were fixed with 4% formaldehyde in PBS at 6 hours post-treatment for 15 minutes at room temperature and then stored in PBS at 4°C. Cells were incubated in blocking buffer (1% FBS and 2% BSA in PBS) and incubated with wheat germ agglutinin conjugated to Alexa Flour 647 (ThermoFisher Scientific, W32466) and a mAb binding to heparan sulfate (AMSBIO Cat# 370255-1, RRID:AB_10891554) at 1:100 dilutions in PBS overnight at 4°C. For co-localization and EFNB1 clustering analysis, EFNB1-FLAG-expressing HPMEC were prepared as described above and transferred to 4°C for 15 min before addition of 10 μg/mL of NS1 and another incubation for 30 min at 4°C. Cells were then either fixed immediately as described above or incubated for 15 min at 37°C before fixation. Cells were permeabilized using 0.2% Triton-X in PBS for 15 min at room temperature and then incubated in blocking buffer for 45 min at room temperature. Primary antibodies targeting the FLAG tag on EFNB1 (Cell Signaling, 8146) and the 6X-histidine tag on NS1 (abcam, ab232492) were added at 1:100 dilution in blocking buffer overnight at 4°C. Cells were washed and then incubated with the fluorescently conjugated secondaries antibodies. All images were acquired using a Zeiss LSM 990 Axio Observer microscope and quantified using Fiji^61^. CellProfiler was used to quantify EFNB1 puncta and clusters using a semi-automated image analysis workflow.

### Bead-based binding assay

Recombinant DENV1-4, ZIKV, and WNV NS1 proteins, as well as the DENV2 NS1 N207Q mutant protein, were commercially acquired from Native Antigen Co. (United Kingdom). Recombinant NS1 domain chimeras and individual NS1 domains were produced in Expi293F cells as described above. Recombinant proteins were randomly biotinylated according to the manufacturer’s instructions (EZ- Link™ Sulfo-NHS-LC-LC-Biotin, No-Weigh™ Format; Thermo Fisher Scientific) and desalted using Zeba Spin Columns (Thermo Fisher Scientific). Recombinant proteins were site biotinylated according to the manufacturer’s instructions (Promega Halotag PEG-Biotin Ligand) and desalted using Zeba Dye and Biotin Removal Spin Columns (Thermo Fisher Scientific). The randomly biotinylated proteins and the site-specifically biotinylated domains were conjugated to avidin-coated MagPlex Luminex beads. Binding complexes were formed in 384-well plates by mixing the appropriately diluted EFNB1 or Gluc (six 4-fold dilutions, starting concentration at 1 μg/mL) with antigen-coupled microspheres for 90 min at 37°C, shaking at 850 rpm. After incubation, plates were washed using an automatic magnetic washer (Tecan Hydrospeed) with PBS containing 0.1% BSA, 0.02% Tween 20 and Proclin 300 (Sigma-Aldrich). Finally, antigen binding was detected using phycoerythrin (PE)-coupled anti-mouse detection antibodies against IgG (Thermo Fisher Scientific). Fluorescence was acquired using an iQue3 (Intellicyt) machine (Sartorius). Binding was extracted as median fluorescence intensity (MFI).

### Enzyme-linked immunosorbent assay

Ninety-six-well plates were coated overnight at 4°C with 5 μg/mL of EFNB1 RBD Fc-fusion proteins in 100 μL of PBS. To minimize non-specific binding, blocking buffer (2% bovine serum albumin (BSA) in PBS) was added for 2 hours at room temperature. After blocking for 1 hour at room temperature, serial dilutions of recombinant DENV NS1, ZIKV NS1, WNV NS1, or Gluc luciferase (as a negative control) were added in a three-fold dilution series (30, 10, 3.33, 1.11, 0.37, and 0.123 μg/mL) in blocking buffer and incubated for 2 hours at 37°C. Plates were washed three times with PBS-T (0.05%), and bound His-tagged recombinant proteins were detected using an anti-His mAb (Abcam, ab232492 at 1:1,000) followed by washing three times with PBS-T. HRP-conjugated secondary antibodies were added for 1 hour at room temperature, avoiding light. Plates were washed five times with PBS-T, and 100 μL of 3,3′,5,5′-Tetramethylbenzidine (TMB) substrate was added and then incubated for 5 to 20 minutes before 50 μL of stop solution (2N H₂SO₄) was added. The microplate was read at 450 nm.

### Dermal vascular leak model

For the murine dermal leak model^33^, the dermis of 6-8-week-old C57BL/6J female mice was shaved and depilated with Nair. After three days, NS1 was mixed with EFNB1-Fc fusion proteins (15 µg each), and the complex or the individual proteins, alongside PBS as a negative control, were intradermally injected into distinct spots (50 µL per spot). The tracer dye 10-kDa dextran conjugated to Alexa Fluor 680 (Sigma; 150 µL, 167 ng/µL) was intravenously injected and allowed to circulate for 3 hours. The dorsal dermis was removed, and extravasated tracer dye was measured using a fluorescence scanner (LI-COR Odyssey CLx Imaging system). The mean fluorescence intensity (MFI) was measured in a consistent circular area surrounding the injection spot using Image Studio software (LI-COR Biosciences).

### Statistical analysis

All quantitative analyses of data were performed and graphed using GraphPad Prism software (v10.5.0). All experiments were repeated at least three times unless indicated. Data are displayed as mean ± SEM. Statistical tests used in this study are indicated in the figure legends, and resulting *P*- values are displayed as * *P* < 0.05, ** *P* < 0.01, and *** *P* < 0.001.

## List of Supplemental Materials

Supplementary Figures 1-4

Supplementary Table 1

## Raw mass spectrometry data

The mass spectrometry-based proteomics data have been deposited at the ProteomeXchange Consortium (http://proteomecentral.proteomexchange.org) via the PRIDE partner repository with the following dataset identifier: PXD067992.

## Acknowledgements

We thank Marco Antonio Chapa and Claudia Sanchez San Martin at UC Berkeley for excellent administrative support. Confocal images were acquired with a Zeiss LSM 990 Axio Observer at the CRL Molecular Imaging Center at UC Berkeley, which is supported by the Gordon and Betty Moore Foundation. We are grateful to Marcus P. Wong, Pedro Carneiro and Sandra Bos for helpful discussions and advice. We thank Kiana D. Dokanchi and Jenny M. Granera-Calero for assistance with animal husbandry.

## Author contributions

Conceptualization: FP, SBB, EH

Methodology: FP, SBB, PRB, PS, EH

Investigation: FP, SRH, CF, XF, SEL, EVJP, JACO, EMD, AHB, NEL, KL, LVT

Visualization: FP, SRH, CF, XF, EVJP, JACO

Funding acquisition: FP, PS, EH

Project administration: PS, EH

Supervision: FP, PS, EH

Writing – original draft: FP, EH

Writing – review & editing: all authors.

## Data and materials availability

Materials generated in this study are available upon request.

## References

1. Paz-Bailey, G., Adams, L. E., Deen, J., Anderson, K. B. & Katzelnick, L. C. Dengue. The Lancet 403, 667–682 (2024).

2. Guzman, M. G. & Harris, E. Dengue. The Lancet 385, 453–465 (2015).

3. Katzelnick, L. C. et al. Antibody-dependent enhancement of severe dengue disease in humans. Science 358, 929–932 (2017).

4. Chen, R. E. et al. Implications of a highly divergent dengue virus strain for cross-neutralization, protection, and vaccine immunity. Cell Host Microbe 29, 1634–1648.e5 (2021).

5. Ooi, E. E. & Kalimuddin, S. Insights into dengue immunity from vaccine trials. Sci. Transl. Med. 15, eadh3067 (2023).

6. Akey, D. L. et al. Flavivirus NS1 Structures Reveal Surfaces for Associations with Membranes and the Immune System. Science 343, 881–885 (2014).

7. Pan, Q. et al. The step-by-step assembly mechanism of secreted flavivirus NS1 tetramer and hexamer captured at atomic resolution. Sci. Adv. 10, eadm8275 (2024).

8. Gutsche, I. et al. Secreted dengue virus nonstructural protein NS1 is an atypical barrel-shaped high-density lipoprotein. Proc. Natl. Acad. Sci. U. S. A. 108, 8003–8008 (2011).

9. Benfrid, S. et al. Dengue virus NS1 protein conveys pro-inflammatory signals by docking onto high-density lipoproteins. EMBO Rep. 1–14 (2022).

10. Biering, S. B. et al. Structural basis for antibody inhibition of flavivirus NS1–triggered endothelial dysfunction. Science 371, 194–200 (2021).

11. Modhiran, N. et al. A broadly protective antibody that targets the flavivirus NS1 protein. Science 371, 190–194 (2021).

12. Wessel, A. W. et al. Human Monoclonal Antibodies against NS1 Protein Protect against Lethal West Nile Virus Infection. mBio 12, e02440–21 (2021).

13. Wessel, A. W. et al. Antibodies targeting epitopes on the cell-surface form of NS1 protect against Zika virus infection during pregnancy. Nat. Commun. 11, 5278 (2020).

14. Yu, L. et al. Monoclonal Antibodies against Zika Virus NS1 Protein Confer Protection via Fc **γ** Receptor-Dependent and -Independent Pathways. mBio 12, e03179–20 (2021).

15. Beatty, P. R. et al. Dengue virus NS1 triggers endothelial permeability and vascular leak that is prevented by NS1 vaccination. Sci. Transl. Med. 7, 1–12 (2015).

16. Libraty, D. H. et al. High circulating levels of the dengue virus nonstructural protein NS1 early in dengue illness correlate with the development of dengue hemorrhagic fever. J. Infect. Dis. 186, 1165–1168 (2002).

17. Wong, M. P. et al. The inflammasome pathway is activated by dengue virus non-structural protein 1 and is protective during dengue virus infection. PLOS Pathog. 20, e1012167 (2023).

18. Chao, C.-H. et al. Dengue virus nonstructural protein 1 activates platelets via Toll-like receptor 4, leading to thrombocytopenia and hemorrhage. PLOS Pathog. 15, e1007625 (2019).

19. Avirutnan, P. et al. Binding of Flavivirus Nonstructural Protein NS1 to C4b Binding Protein Modulates Complement Activation. J. Immunol. 187, 424–433 (2011).

20. Chung, K. M. et al. West Nile virus nonstructural protein NS1 inhibits complement activation by binding the regulatory protein factor H. Proc. Natl. Acad. Sci. 103, 19111–19116 (2006).

21. Avirutnan, P. et al. Antagonism of the complement component C4 by flavivirus nonstructural protein NS1. J. Exp. Med. 207, 793–806 (2010).

22. Wang, C. et al. Endocytosis of flavivirus NS1 is required for NS1-mediated endothelial hyperpermeability and is abolished by a single N-glycosylation site mutation. PLOS Pathog. 15, 1–33 (2019).

23. Puerta-Guardo, H. et al. Zika Virus Nonstructural Protein 1 Disrupts Glycosaminoglycans and Causes Permeability in Developing Human Placentas. J. Infect. Dis. 221, 313–324 (2020).

24. Puerta-Guardo, H., Glasner, D. R. & Harris, E. Dengue Virus NS1 Disrupts the Endothelial Glycocalyx, Leading to Hyperpermeability. PLoS Pathog. 12, 1–29 (2016).

25. Puerta-Guardo, H. et al. Flavivirus NS1 Triggers Tissue-Specific Disassembly of Intercellular Junctions Leading to Barrier Dysfunction and Vascular Leak in a GSK-3β-Dependent Manner. Pathogens 11, 615 (2022).

26. Puerta-Guardo, H. et al. Flavivirus NS1 Triggers Tissue-Specific Vascular Endothelial Dysfunction Reflecting Disease Tropism. Cell Rep. 26, 1598–1613.e8 (2019).

27. Lo, N. T. N. et al. Molecular Determinants of Tissue Specificity of Flavivirus Nonstructural Protein 1 Interaction with Endothelial Cells. J. Virol. 96, e0066122 (2022).

28. Laing, E. D. et al. Structural and functional analyses reveal promiscuous and species specific use of ephrin receptors by Cedar virus. Proc. Natl. Acad. Sci. 116, 20707–20715 (2019).

29. Pryce, R. et al. A key region of molecular specificity orchestrates unique ephrin-B1 utilization by Cedar virus. Life Sci. Alliance 3, e201900578 (2020).

30. Negrete, O. A. et al. EphrinB2 is the entry receptor for Nipah virus, an emergent deadly paramyxovirus. Nature 436, 401–405 (2005).

31. Pernet, O., Wang, Y. E. & Lee, B. Henipavirus Receptor Usage and Tropism. in Henipavirus (eds Lee, B. & Rota, P. A.) vol. 359 59–78 (Springer Berlin Heidelberg, Berlin, Heidelberg, 2012).

32. Pahmeier, F. et al. Antibodies targeting Crimean-Congo hemorrhagic fever virus GP38 limit vascular leak and viral spread. Sci. Transl. Med. 17, eadq5928 (2025).

33. Glasner, D. R. et al. Dengue virus NS1 cytokine-independent vascular leak is dependent on endothelial glycocalyx components. PLoS Pathog. 13, 1–22 (2017).

34. Sato, S., et al. EPHB2 carried on small extracellular vesicles induces tumor angiogenesis via activation of ephrin reverse signaling. JCI Insight 4, e132447 (2019).

35. 35. Passaro, S. et al. Boltz-2: Towards Accurate and Efficient Binding Affinity Prediction. Preprint at 10.1101/2025.06.14.659707 (2025).

36. Lee, H.-S., Nishanian, T. G., Mood, K., Bong, Y.-S. & Daar, I. O. EphrinB1 controls cell–cell junctions through the Par polarity complex. Nat. Cell Biol. 10, 979–986 (2008).

37. Park, I. & Lee, H.-S. EphB/ephrinB Signaling in Cell Adhesion and Migration. Mol. Cells 38, 14–19 (2015).

38. Daar, I. O. Non-SH2/PDZ reverse signaling by ephrins. Semin. Cell Dev. Biol. 23, 65–74 (2012).

39. Gschwendt, M. et al. Inhibition of protein kinase C μ by various inhibitors. Inhibition from protein kinase c isoenzymes. FEBS Lett. 392, 77–80 (1996).

40. Hafirassou, M. L. et al. A Global Interactome Map of the Dengue Virus NS1 Identifies Virus Restriction and Dependency Host Factors. Cell Rep. 21, 3900–3913 (2017).

41. Scaturro, P. et al. An orthogonal proteomic survey uncovers novel Zika virus host factors. Nature 561, 253–257 (2018).

42. Shah, P. S. et al. Comparative Flavivirus-Host Protein Interaction Mapping Reveals Mechanisms of Dengue and Zika Virus Pathogenesis. Cell 175, 1931–1945.e18 (2018).

43. Barbachano-Guerrero, A., Endy, T. P. & King, C. A. Dengue virus non-structural protein 1 activates the p38 MAPK pathway to decrease barrier integrity in primary human endothelial cells. J. Gen. Virol. 101, 484–496 (2020).

44. Tanaka, M., Kamata, R. & Sakai, R. Phosphorylation of ephrin-B1 via the interaction with claudin following cell–cell contact formation. EMBO J. 24, 3700–3711 (2005).

45. Modhiran, N. et al. Dengue virus NS1 protein activates cells via Toll-like receptor 4 and disrupts endothelial cell monolayer integrity. Sci. Transl. Med. 7, 1–11 (2015).

46. Alcalá, A. C., et al. The dengue virus non-structural protein 1 (NS1) uses the scavenger receptor B1 as a cell receptor in cultured cells. J. Virol. 96, e0166421 (2022).

47. Davy, A., Aubin, J. & Soriano, P. Ephrin-B1 forward and reverse signaling are required during mouse development. Genes Dev. 18, 572–583 (2004).

48. Huynh-Do, U. et al. Ephrin-B1 transduces signals to activate integrin-mediated migration,attachment and angiogenesis. J. Cell Sci. 115, 3073–3081 (2002).

49. Hanley, J. P. et al. Immunotranscriptomic profiling the acute and clearance phases of a human challenge dengue virus serotype 2 infection model. Nat. Commun. 12, 3054 (2021).

50. Waickman, A. T. et al. Low-dose dengue virus 3 human challenge model: a phase 1 open-label study. Nat. Microbiol. 9, 1356–1367 (2024).

51. Banerjee, A. et al. RNA-Seq analysis of peripheral blood mononuclear cells reveals unique transcriptional signatures associated with disease progression in dengue patients. Transl. Res. 186, 62–78.e9 (2017).

52. Woodruff, T. M. et al. Epha4-Fc Treatment Reduces Ischemia/Reperfusion-Induced Intestinal Injury by Inhibiting Vascular Permeability. Shock 45, 184–191 (2016).

53. Khan, N. et al. Inhibiting Eph/ephrin signaling reduces vascular leak and endothelial cell dysfunction in mice with sepsis. Sci. Transl. Med. 16, eadg5768 (2024).

54. Cercone, M. A., Schroeder, W., Schomberg, S. & Carpenter, T. C. EphA2 receptor mediates increased vascular permeability in lung injury due to viral infection and hypoxia. Am. J. Physiol.- Lung Cell. Mol. Physiol. 297, L856–L863 (2009).

55. Krump-Konvalinkova, V. et al. Generation of Human Pulmonary Microvascular Endothelial Cell Lines. Lab. Invest. 81, 1717–1727 (2001).

56. Yang, X. et al. A public genome-scale lentiviral expression library of human ORFs. Nat. Methods 8, 659–661 (2011).

57. Biering, S. B. et al. SARS-CoV-2 Spike triggers barrier dysfunction and vascular leak via integrins and TGF-β signaling. Nat. Commun. 13, 7630 (2022).

58. Sanjana, N. E., Shalem, O. & Zhang, F. Improved vectors and genome-wide libraries for CRISPR screening. Nat. Methods 11, 783–784 (2014).

59. Pahmeier, F. et al. Identification of host dependency factors involved in SARS-CoV-2 replication organelle formation through proteomics and ultrastructural analysis. J. Virol. 97, e00878–23 (2023).

60. Scaturro, P. et al. An orthogonal proteomic survey uncovers novel Zika virus host factors. Nature 561, 253–257 (2018).

61. Schindelin, J. et al. Fiji: An open-source platform for biological-image analysis. Nat. Methods 9, 676–682 (2012).

