## Supplementary Figures 1-4 for "Dengue Virus NS1 Binds Ephrin B1 to Trigger Endothelial Dysfunction"

**Figures S1-S4**

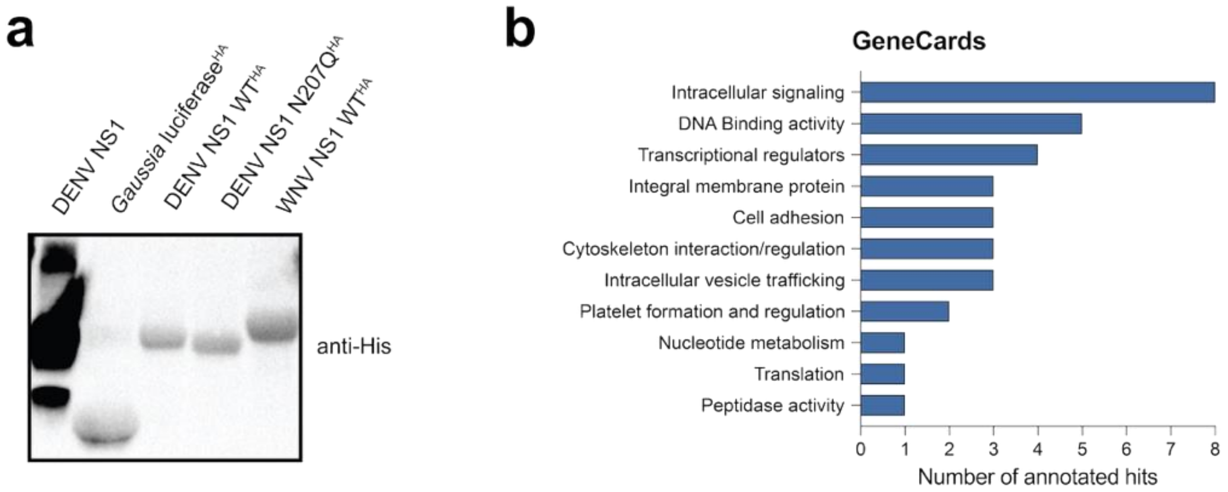


**Figure S1**. **Recombinant protein validation and GeneCards analysis of host interaction partners**. **a**, Recombinant HA- and His-tagged proteins were analyzed by Western blot using anti-His antibodies. **b,** GeneCard annotations of DENV NS1 WT-selective host factors.

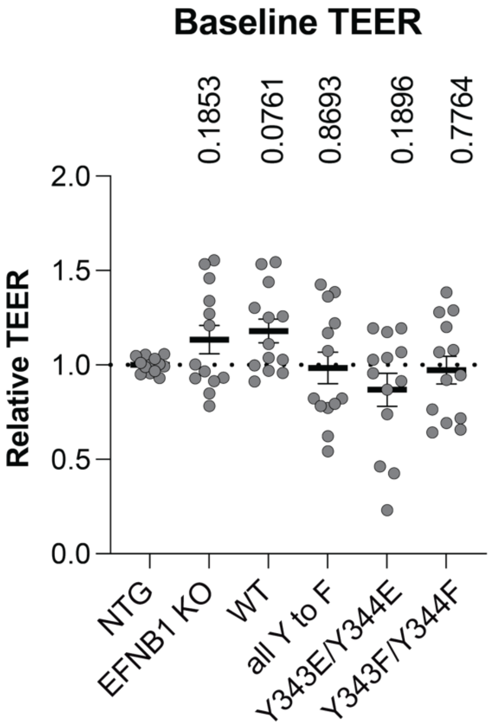


**Figure S2. Genetically modified HPMEC cell lines form endothelial barriers *in vitro*.** TEER values at 0 and 24 hours post-treatment of the HPMEC EFNB1-KO cell line transduced with EFNB1 expression cassettes. Statistical comparisons were performed by ordinary one-way ANOVA and *P*-values are shown on the top of each dataset. NTG, non-targeting guide; KO, knockout.

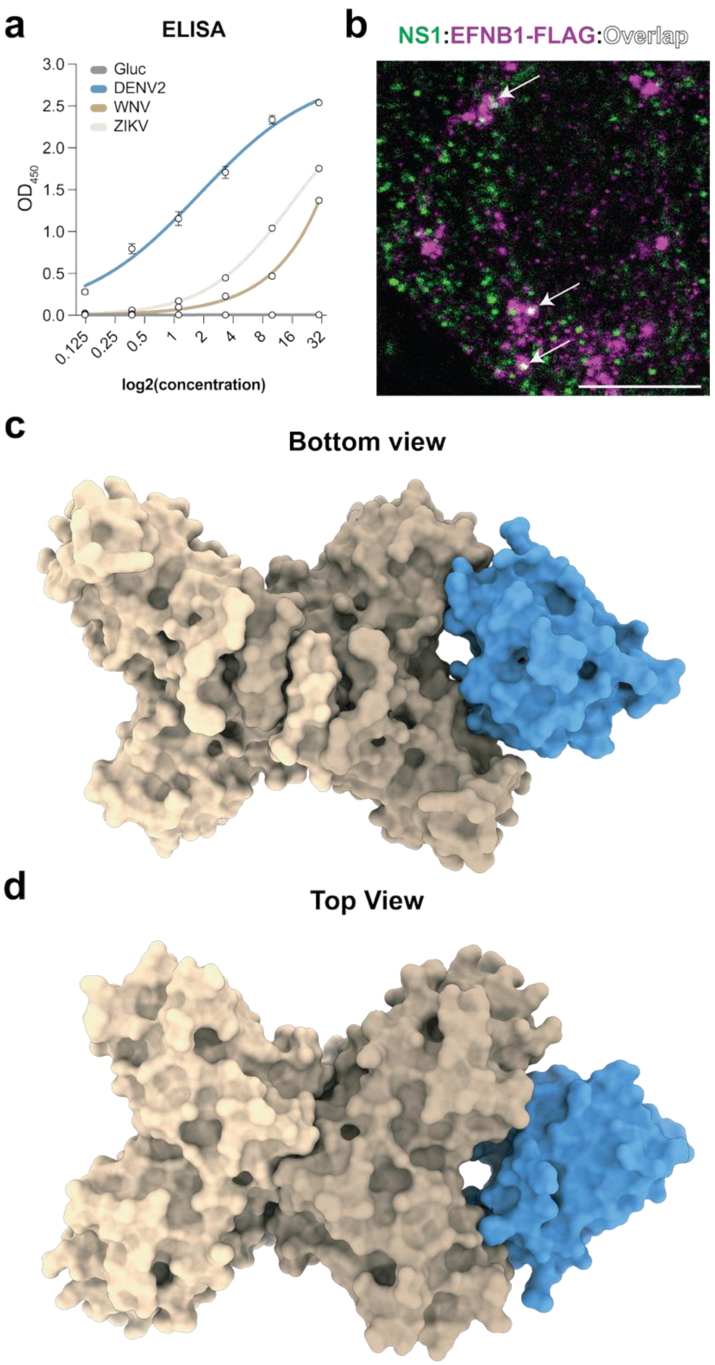


**Figure S3. EFNB1 binds and co-localizes with NS1. a**, Binding of flavivirus NS1 proteins to EFNB1-Fc fusion in an ELISA. Plates were coated with the EFNB1-RBD Fc fusion protein, a dilution series of the indicated His-tagged NS1 proteins was added, and their binding was detected using an anti-His antibody (n=2). **b**, EFNB1-FLAG-expressing cells were treated with NS1 or left untreated and incubated for 30 minutes at 4°C and then fixed. DENV NS1 (green) and EFNB1-FLAG (purple) localization was detected with mAbs targeting the His or FLAG tag, respectively (scale bar, 10 μm). **c-d**, Surface representation of EFNB1 RBD (blue) interacting with dimeric DENV2 NS1 (tan) as predicted by Boltz2.

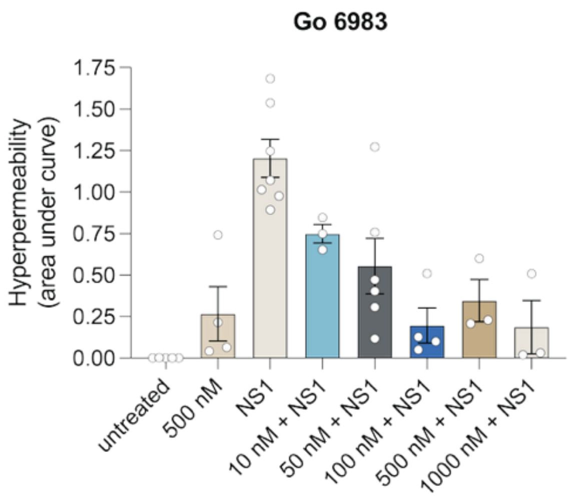


**Figure S4. PKC inhibition can block NS1-mediated endothelial dysfunction.** HPMECs were treated with NS1 in absence of decreasing concentrations (in nM) of the PKC inhibitor Go 6983, and the electrical resistance was measured at 0, 6 and 24 hours post-treatment. The area under the curve was plotted as the mean ± SEM (n≥2).
